# Evaluation of the gene fusion landscape in early onset sporadic rectal cancer reveals association with chromatin architecture and genome stability

**DOI:** 10.1101/2024.04.09.588655

**Authors:** Asmita Gupta, Sumedha Avadhanula, Murali Dharan Bashyam

## Abstract

Gene fusions represent a distinct class of structural variants identified frequently in cancer genomes across cancer types. Several gene fusions exhibit gain of oncogenic function and thus have been the focus of development of efficient targeted therapies. However, investigation of fusion landscape in early-onset sporadic rectal cancer, a poorly studied colorectal cancer subtype prevalent in developing countries, has not been performed. Here, we present a comprehensive landscape of gene fusions in EOSRC and CRC using patient derived tumor samples and data from The Cancer Genome Atlas, respectively. Gene Ontology analysis revealed enrichment of unique biological process terms associated with 5’- and 3’- fusion partner genes. Extensive network analysis highlighted genes exhibiting significant promiscuity in fusion formation and their association with chromosome fragile sites. Investigation of fusion formation in the context of global chromatin architecture unravelled a novel mode of gene activation that arose from fusion between genes located in orthogonal chromatin compartments. The study provides novel evidence linking fusions to genome stability and architecture and unearthed a hitherto unidentified mode of gene activation in cancer.

## Introduction

Genomic instability, a hallmark of cancer^1^, manifests in various forms of aberrant chromosomal rearrangements including translocations^2,3^, inversions^4^ and tandem genomic insertions^5^, that may result in Gene Fusions (GFs). The Philadelphia chromosome^6^, resulting from translocation between chromosomes 9 and 22 (t(9;22)), and leading to the *BCR-ABL1* fusion in chronic myeloid leukaemia (CML)^7^, was the first GF to be studied. This GF has been the focus for developing several therapies targeted against the tyrosine kinase ABL1, including the first generation tyrosine kinase inhibitor (TKI) imatinib^8^, followed by second and third generation TKIs like nilotinib^9^ and ponatinib^10^. Subsequent work revealed frequent occurrences of GFs in several haematological malignancies e.g. fusions involving immunoglobulins (*IgH/K-MYC* in Burkitt lymphoma^11^ and *IgH/K-BCL2/BCL6/MYC* in diffuse large B-cell lymphoma^12^), *PML-RARA* in acute promyelocytic leukemia^13^ and *ETV6-RUNX1* in acute lymphoblastic leukemia^14^. These GFs were validated as the major chromosomal anomalies in early leukemogenesis^12,15^. GFs have also been studied across several solid tumour types, including the highly recurrent *TMPRSS2-ERG* fusion in prostate cancer (∼90% of all prostate cancers tested clinically)^16^, that has been comprehensively validated for diagnostic utility^17^. Similarly, the *EML4-ALK1* fusion identified in non-small cell lung cancer has been the focus of therapeutic approaches targeting the tyrosine kinase ALK1 such as crizotinib^18^. A recent Pan-cancer analysis revealed the presence of several cancer-type specific GFs and prevalence of kinase proteins in the fusion partners^19^, supporting the prevalence and clinical utility of GFs in neoplasms.

Among solid tumours, colorectal cancer (CRC) constitutes the fourth most prevalent cancer type in terms of incidence and third in terms of mortality (GLOBOCAN 2020, (http://gco.iarc.fr); age standardized rates, both sexes). Due to the identification and removal of early lesions through efficient implementation of screening modalities, CRC incidence is receding in the West, though death rates remain high^20,21^. In contrast, there is an alarming worldwide increase in incidence rates of rectal cancer in the young (GLOBOCAN 2020; cancer incidence in five continents (https://ci5.iarc.fr)); early-onset sporadic rectal cancer (EOSRC) is the predominant CRC subtype in the developing world including India^22^. Earlier studies have characterized GFs in primary and metastatic CRC^23,24^, and GFs involving actionable cancer driver genes such as *NTRK1/2/3, RET, ALK, BRAF* and *FGFR2* were identified^23–25^. However, no studies evaluating GFs in EOSRC have been performed till date.

Chromosome fragile sites (CFS) act as genomic fault lines vulnerable to breaks under replicative stress, and have been reported as potential hotspots for aberrant translocation and recombination^26,27^. However, no systematic investigation to correlate CFS with GF breakpoints has been undertaken till date. Secondly, genomic rearrangements can also be modulated by spatial chromatin organization^28^, another facet of GF formation yet to be explored in detail.

Here, we define the landscape of GFs in early-onset sporadic rectal cancer (EOSRC), based on RNA sequencing data generated from patient-derived tumour tissues, and describe the molecular features of genes participating in GFs. We report an extensive network formed by ‘promiscuous’ genes which formed GFs with multiple fusion partners. We further report a significant correlation between recurrent GF breakpoints arising from these promiscuous genes and otherwise, and CFSs. Finally, our results reveal a novel mode of gene activation of the 3’- fusion partner in GFs formed between 5’ and 3’ partners located in open (‘A’) and close (‘B’) chromatin compartments, respectively.

## Methods

### Sample collection and dataset features

GF identification was performed on 37 EOSRC samples (**Figures 1a and S1a** and **Table S1),** a part of a well annotated microsatellite stable CRC tissue repository built during the past two decades^22,29,30^. For a comparative analysis of GFs in CRC, we accessed the paired-end, raw transcriptomics data for rectal (TCGA-READ) and colon (TCGA-COAD) adenocarcinoma samples from The Cancer Genomic Atlas (TCGA) Genomic Data Commons (GDC) legacy archive (https://portal.gdc.cancer.gov/). Data from samples with ambiguous annotation status were not included. The samples selected for the study represented primary adenocarcinomas from patients with no prior treatment or family history or secondary malignancies. Based on these criteria, we accessed RNA sequencing data from 55 TCGA-READ and 102 TCGA-COAD samples from the GDC portal. For most analyses, we pooled TCGA-COAD and TCGA-READ samples (referred to as TCGA-CRC), unless otherwise stated. Additional GF identification and analysis was carried out on raw transcriptome sequencing data of five well-established colorectal cancer cell-lines viz. Caco2, HCT116, HT29, SW480, and SW620, as well as three additional cell lines viz. MCF-7, PC3 and PANC-1 obtained from the Cancer Cell Line Encyclopaedia project (CCLE) project. The data were downloaded from the Gene Expression Omnibus (GEO) server with following accession ID – PRJNA523380.

**Figure 1:**
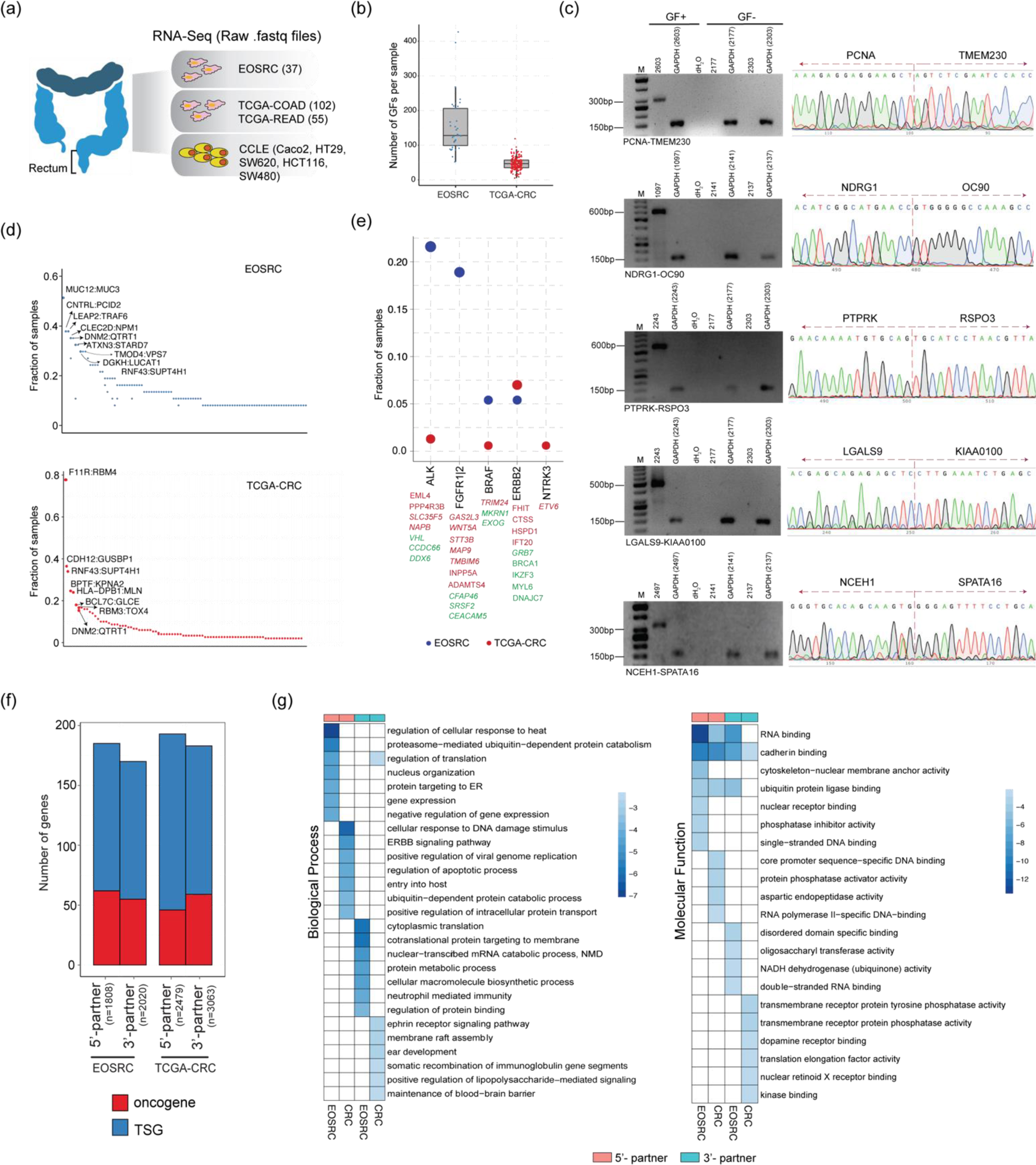
(a) Schematic describing different datasets used for GF identification. (b) Distribution of per sample GFs in EOSRC and TCGA-CRC. (c) RT-PCR validation of GFs identified in EOSRC. GF+ and GF-represent sample/s with or without fusion, respectively. M, 50bp DNA ladder (Invitrogen, Carlsbad, CA, USA). The vertical broken line above the electropherograms indicate the breakpoint junction between the two GF partner genes. (d) Measure of GF recurrence in EOSRC and CRC; GFs identified in >=10, and >=25 samples for EOSRC and CRC, respectively, are labelled. (e) Recurrence of known potentially actionable genes participating in GFs. The corresponding 5’- and 3’- fusion partners of the genes on the x-axis are indicated in red and green, respectively. GF partners reported in this work are shown in italics. (f) Proportion of annotated oncogenes and tumour suppressors participating in GF formation. (g) GO enrichment of 5’- and 3’- partner genes participating in GF formation. The color scale indicates log10 (p-values) associated with the terms.

### RNA-Sequencing for EOSRC samples

Total RNA from tissue samples was isolated using the TriZol reagent (Life Technologies, Grand Island, NY, USA) and RNeasy Kit (Qiagen, Valencia, CA, USA) following the manufacturer’s protocol. Transcriptome sequencing was outsourced to Medgenome Labs Ltd, Bangalore, India. The libraries were prepared using Illumina TruSeq2 kit, following rRNA depletion and sequenced on the HiSeq 2000 to generate 2×100 bp paired-end sequence reads with sequencing depth ranging from 140-200 million reads per sample. Both raw sequencing data in the form of *fastq* files and processed data in the form of transcript counts are available on Gene Expression Omnibus (GEO) database **(GSE253106).**

### Detection of putative GFs

Quality assessment of raw EOSRC RNA-Seq data was done using FastQC (https://www.bioinformatics.babraham.ac.uk/projects/fastqc/) and subsequent filtering was done using Trimmomatic^31^. Reads with a Phred quality score of <30 and length <35 were discarded. For EOSRC, three computational tools viz. FusionCatcher (FC), SOAPFuse (SF), and Arriba were used with their default parameters. Experimentally verified false positive GFs identified from the ConjoinG database (https://metasystems.riken.jp/conjoing/index), GTEx database^32^, as well as from published literature^33^ were removed (**Figure S1b**). GF calls where both the partner genes were the same, and where a sample consisted of duplicate calls, as determined by identical breakpoint in a single sample, were discarded. GFs between homologous genes or pseudogenes were also removed. A non-redundant list of GFs, pooled from calls made by each algorithm was used for all subsequent *in-silico* analyses, to ensure integration of complementary features of each algorithm.

For the TCGA-CRC data set, GFs were detected using FusionCatcher and Arriba (we avoided SOAPFuse since it requires relatively longer computational time, making it difficult to analyze large number of samples) with the same set of filters as applied for previous sets. GFs were detected from transcriptome data of cell lines using FusionCatcher and Arriba with the same methodology. For the pan-cancer analysis, GFs detected using Arriba and STAR-Fusion were downloaded from the TCGA portal (https://portal.gdc.cancer.gov/) for 31 different cancer types (**Table S1)**. The same set of filters as described above were used for filtering out false positive GF calls, to obtain a final list of GFs for further analysis.

### GF validation using Reverse Transcription – Polymerase Chain Reaction (RT-PCR) followed by Sanger sequencing

EOSRC GFs identified by at least two algorithms and exhibiting a spanning read support of >5 and junction read support of >3 were selected for validation by RT-PCR. The strategy for PCR primer design is depicted in **Figure S1c**. The GF breakpoints identified by *in-silico* tools were confirmed using Sanger sequencing as per standard protocol.

### Analysis of fusion partner genes

For a categorical classification of fusion partner genes, the Cancer gene census (CGC) dataset was obtained from the COSMIC database^34^ (cancer.sanger.ac.uk; updated as of May 7, 2022); dataset of tumour suppressor genes was downloaded from Tumor Suppressor Gene Database (TSGene)^35^; and list of kinases was obtained from published literature^19^. Functional designation of 5’- and 3’- fusion partner genes based on Gene Ontology (GO), was performed using Enrichr^36^ and redundant GO terms were summarized using Revigo^37^. The GF network was created and analyzed using igraph library from R (https://github.com/igraph/rigraph) and visualized using Cytoscape^38^. The network nodes were characterized in terms of metrics including the *degree* and *betweenness/centrality* scores as quantified by igraph. The degree measures the number of GFs. Betweenness (or centrality) is a measure of how often a node occurs on all shortest paths between two nodes.

### Analyses of chromosomal fragile sites (CFS)

For estimating overlap between the GF breakpoints and CFSs, we used publicly available HCT116 FANCD2 ChIP-Seq data (GEO accession – GSE141101), and the U2OS Mitotic DNA Synthesis – followed by sequencing (MDS) data (GEO accession – GSE149376), to reliably map high confidence CFS regions. A description of these datasets including their GEO accessions is provided in **Table S2**. Briefly, raw sequencing data from FANCD2 ChIP-Seq was subjected to sequence quality assessment using FastqQC and filtering using Trimmomatic^31^, and reads were mapped to GRCh38 reference assembly of the human genome using bowtie2^39^. Peaks were calculated using MACS2^40^ and bedtools^41^ was used to estimate overlap of GF breakpoints with resulting BED files. For MDS data, the processed peak files in BED format were directly accessed from the GEO accession for subsequent analysis.

### Analyses of Hi-C data

Raw Hi-C sequencing data for HCT116 and SW480 CRC cell lines were obtained from GEO bioproject PRJNA553150^42^ and for MCF-7, PC3, and PANC-1 cell lines from PRJNA695073^43^, PRJNA297887^44^, and PRJNA767590^45^, respectively. The data were processed using HiC-Pro^46^ and normalized contact matrices were obtained at 40Kb, 60Kb and 1Mb resolutions, separately. Replicates were pooled for each cell line and valid contact pairs were estimated. Contact matrices at all resolutions were balanced using the Iterative Correction and Eigenvector decomposition (ICE)^47^ method, implemented in the HiC-Pro^46^ pipeline. FitHiC2^48^ was used to estimate the statistically significant intra- and inter-chromosomal contacts from ICE normalized contact matrices at 40Kb resolution and observed contacts with q-values < 0.05 were considered for contact estimation. HiGlass^49^ was used for visualization of contact matrices. The contribution of each chromosome towards GF formation was estimated by normalizing the number of GFs by respective chromosome length to remove any bias coming from differing chromosome lengths. Chromatin compartments were analysed through the eigenvector analysis of the Pearson correlation matrix at 40Kb resolution, obtained from the HiTC^50^ package, with ‘open’ and ‘close’ compartments indicated by ‘A’ and ‘B’ nomenclature, respectively.

### Transcript quantification from RNA-Seq data and quantitative RT-PCR (RT-qPCR)

After quality assessment and filtering as described above, reads were aligned to GRCh38 reference human genome assembly from Ensemble using STAR^51^ and transcript quantification was performed using featureCounts^52^ and RSEM^53^. RT-qPCR was performed as described earlier^54^.

## Results

### Overview of molecular features of GF partner genes

Based on a rigorous analysis pipeline displayed in **Figure S1b** and described in materials and methods, we identified 3062 GFs (median per sample GF of 128) across 37 EOSRC samples (**Figures 1b and S2a, Table S1;** complete list of EOSRC GFs is in **Table S3)**. Similarly, we identified 4150 GFs (median per sample GF of 47) from 152 TCGA-CRC samples (**Figures 1b and S2a**). A corresponding TCGA pan cancer analysis revealed the spectrum of sample-wise GF frequency across multiple cancer types highlighting a higher frequency in TCGA-OV/STAD/GBM/ESCA compared to other cancer types (**Figure S2b).** Here, the median GFs in the TCGA-COAD and READ datasets were estimated to be 27 and 38, respectively (**Figure S2b, inset**), similar to our estimate for TCGA-CRC. A comparison of GF calls made by the three approaches for EOSRC and two approaches for TCGA-CRC and five CRC cell lines revealed poor concordance **(Figure S2c-e)**, highlighting the inherent differences between various GF predicting approaches and underpinning the importance of using multiple pipelines. Further, we validated a significant fraction of the GFs identified in EOSRC by RT-PCR and confirmed their predicted breakpoints by Sanger sequencing (**Figures 1c and S3a**).

We identified several recurrent GFs in EOSRC namely *MUC12-MUC3A* (54%, 20 samples), *CNTRL-PCID2* and *LEAP2-TRAF6* (37%, 14 samples each), and *CLEC2D-NPM1* and *DNM2-QTRT1* (35%, 13 samples) (**Figure 1d**). Of these, the first three are not reported earlier underscoring the importance of our attempt to evaluate EOSRC, a hitherto poorly studied CRC subtype. Similarly, the recurrent GFs in TCGA-CRC included *F11R-RBM4* (76%; 117 samples), *GUSBP1-CDH12* (33%, 52 samples), *RNF43-SUPT4H1* (33%, 51 samples), *BPTF-KPNA2* and *HLA-DPB1-MLN* (37 and 36 samples (24% each), respectively). All, except *RNF43-SUPT4H1*, are not reported earlier. Notably, *DNM2-QTRT1* (35% in EOSRC, 15.7% in TCGA-CRC) and *RNF43-SUPT4H1* (27% in EOSRC, 33% in CRC) were recurrently observed in both datasets (**Figure 1d**). Novel GFs involving potentially actionable genes *FGFR1/2* and *ALK*^23^ were observed in EOSRC, and involving *NTRK3*^25^ in TCGA-CRC (**Figure 1e**). Further, previously well-characterized GFs *PTPRK-RSPO3* and *EIF3E-RSPO2*^55^ were detected in both EOSRC and TCGA-CRC, while *TCF7L2-VT11A*^56^ was detected only in EOSRC (**Figure S3b)**.

We next checked whether a specific category of genes was enriched in the GFs identified in this study. Given that majority of cancer associated genes are tumor suppressors (Cancer Gene Census database^57^), we detected a higher representation of tumour suppressors (compared to oncogenes) in both 5’ and 3’ fusion partners (**Figure 1f**) in EOSRC as well as TCGA-CRC whereas, no significant difference in enrichment or depletion of either oncogenes or tumour suppressors was evident between 5’- and 3’- genes. Furthermore, GO analysis revealed enrichment of distinct ‘Biological Processes’ terms, with essentially no overlap between 5’ and 3’ partners or between the EOSRC and TCGA-CRC data sets (**Figure 1g**). However, there was significant enrichment of GO ‘Molecular Function’ terms RNA binding, Cadherin binding, and Ubiquitin protein ligase binding in the 5’ partners of both data sets. In addition, Cadherin binding was also enriched in the 3’-partners of both data sets. Furthermore, while the ‘Molecular Function’ term Disordered domain binding was enriched in 3’-partners in EOSRC, Kinase binding and Receptor protein activity were enriched in 3’-partners in TCGA-CRC (**Figure 1g**). Enrichment of similar biological pathways was also detected in a previous study on chimeric RNAs in CRC^58^. Finally, we investigated the fraction of 5’- and 3’- partner genes forming unique or multiple GFs with other genes. A majority of genes (81% and 65% of 5’- genes, and 90% and 86% of 3’- genes in EOSRC and TCGA-CRC, respectively) were involved in unique GF pairs. Moreover, while 17% and 35% of 5’- partners in EOSRC and TCGA-CRC, respectively, participated in GF formation with ≥ 2 different genes; 8.7% and 13.7% of 3’- genes in EOSRC and TCGA-CRC, respectively, showed promiscuity in GF formation (**Figure S3c**). Thus, a relatively higher fraction of 5’- genes appeared to function as promiscuous gene partners in GFs compared to 3’- genes.

### Promiscuous fusion partners form extensive gene networks

The identification of genes participating in multiple GFs prompted us to investigate GF networks^59,60^. We identified a central large network with extended sub-networks at the periphery, the latter connected to the central large network via smaller nodes in both EOSRC (**Figures 2a and S4a)** and TCGA-CRC (**Figure S5a**). Additionally, both EOSRC and TCGA-CRC had smaller independent networks which were mostly centered around single genes (**Figures S4b-g and S5b-e**). The single large GF network in EOSRC was centered around *EEF1A1, IgH, IgK*, *CEACAM5*, and *PTMA,* all of which displayed highest degrees and centrality scores (**Figure 2b**); and connected to multiple peripheral networks centered around *WWOX, KRT8, RN7SL1, MUC2, PIGR, NPM1, FGFR2, ACTB, WDR74*, *TMPO* and *NEAT1* (**Figure 2a**). The largest GF network in TCGA-CRC was defined by *EEF1A1, PIGR, CEACAM5, PDIA3*, and *PTMA*, which also exhibited highest degrees and centrality scores (**Figure 2b**), while *ACTB, SELENBP, Y_RNA, ERBB2* and *KRT8* formed the peripheral networks (**Figure S6**).Thus, three genes (*EEF1A1, PTMA*, and *CEACAM5*) were common to EOSRC and TCGA-CRC in contributing towards the single large network. Genes exhibiting high degree and centrality in EOSRC and TCGA-CRC are enumerated in **Figure S7a**. A closer inspection of the central networks in EOSRC and TCGA-CRC revealed that a few genes with high centrality scores acted as connectors between the major central nodes (with degree ≥ 10) and the peripheral nodes (**Figure 2b**). For example, *KRT8* connected the nodes formed by *IgH* and *IgK* to *MUC2* in EOSRC (**Figure 2a**). *KRT8* was also linked to a hub created by *ACTB*, which in turn, was further connected to *CEACAM5*. Similarly, *WDR74* connected the central *EEF1A1* node to a peripheral node formed by *PIGR*. On the other hand, in TCGA-CRC, we identified an extensive *ERBB2* mediated cluster of GFs, which was connected to the single large network through *EEF1A1* **(Figure S6)**. In addition, *KRT8* connected a *MUC2* mediated sub-network to major *PDIA3* and *CEACAM5* nodes besides forming link with the peripheral subnetwork centred around *ACTB* (**Figure S6)**. Again, it was interesting to observe *MUC2* and *KRT8* as ‘connectors’ in both data sets.

**Figure 2:**
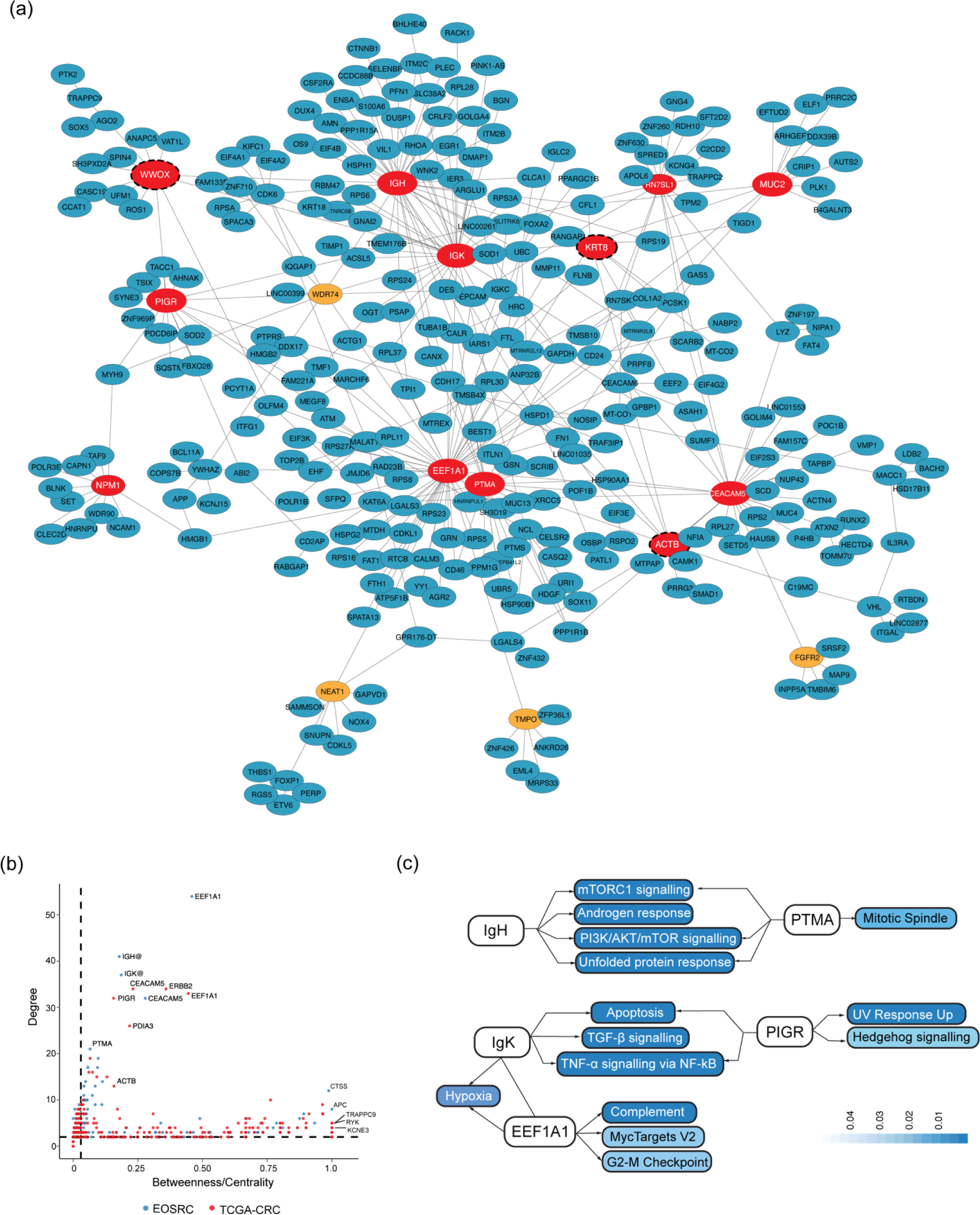
GF network in EOSRC. (a) The largest network filtered to include nodes with a total degree > 3 and betweenness > 0.1. Each node/vertex represents a gene, while edge/connection represents fusion between the genes. The nodes highlighted in red and orange have degree ≥ 10, and 5 ≤ degree < 10, respectively. Singletons and nodes connected by single links to another node are not shown. Nodes with broken outline indicate genes associated with CFS loci (b) Distribution of degree and betweenness/centrality metrics for EOSRC and TCGA-CRC GF networks. Broken vertical and horizontal lines indicate degree and betweenness/centrality values of 1 and 0.05, respectively. (c) MSigDB hallmark pathway enrichment of genes forming fusions with *IgH*, *IgK*, *EEF1A1*, *PTMA*, and *PIGR* in EOSRC. The pathways are indicated in colored boxes with color scale indicating the p-values (Fisher’s exact test) of extent of pathway enrichment. Links without arrowheads represent fusion between genes (*IgK* and *EEF1A1*), while links with arrowheads indicate association of partner genes with the pathways.

The network analysis led us to investigate the possible enrichment of cancer specific pathways and functional categories in genes which formed fusions with central node genes in EOSRC. Indeed, a pathway enrichment analysis using MSigDB hallmark gene sets revealed that while *IgH* and *PTMA* preferentially formed fusions with genes involved in mTORC1 signalling, PI3K/AKT pathway, and regulation of spindle formation, *IgK* and *PIGR* formed fusions with genes involved in critical cancer related pathways like apoptosis and TNF-alpha signalling via NF-kB in EOSRC (**Figure 2c, Figure S7b**). A parallel GO analysis suggested that fusion partners of node genes were associated with distinct and cancer related biological pathways such as response to oxidative stress, regulation of cell differentiation, and apoptotic processes (**Figure S7c**).

Interestingly, three of ten genes with highest degree in the GF network, *WWOX*, *ACTB*, and *KRT8* (**Figure S7a**), overlapped with well-annotated CFS loci FRA16D^61^, FRA12A^62^, and FRA7B^62^, respectively. This suggested a possible role of CFS in driving promiscuity in GF partners. Hence, we proceeded to evaluate correlation between GF breakpoints of genes forming fusion with multiple partners and CFSs.

### GF breakpoints for promiscuous genes overlap with CFS

We used publicly available datasets for FANCD2 ChIP-Seq and MDS (**Table S2**), to document genome wide CFSs. Of all genes which formed GFs in EOSRC (either 5’ or 3’- fusion partners), ∼5% harbored breakpoints overlapping with CFS. Interestingly, this fraction consistently increased for genes which formed GFs with ≥2 (p-value = 0.06, Fisher’s exact test) and >5 partners (p-value <0.05, Fisher’s exact test) partners respectively (**Figure 3a**). This trend was also observed in the TCGA-CRC dataset (**Figure 3a**), suggesting that the breakpoints in promiscuous genes which form GFs with multiple partners, might have a higher probability of arising from a CFS loci. Unlike the trend observed for promiscuous genes, we did not find any significant association between sample-wide recurrent GF breakpoints (for both 5’- and 3’- partners) and CFS enrichment (**Figure S8a**). A comprehensive analysis of all GF breakpoints (either 5’ or 3’) which overlapped with CFS loci (either FANCD2 or MDS) revealed that the fraction of inter- and intra-chromosomal GFs overlapping with CFS loci were similar in EOSRC (49% and 52% respectively) (**Figure S8b**). On the other hand, the overlap of CFSs with intra-chromosomal GFs (76%) was significantly higher than with inter-chromosomal GFs (24%) in TCGA-CRC (**Figure S8b**).

**Figure 3:**
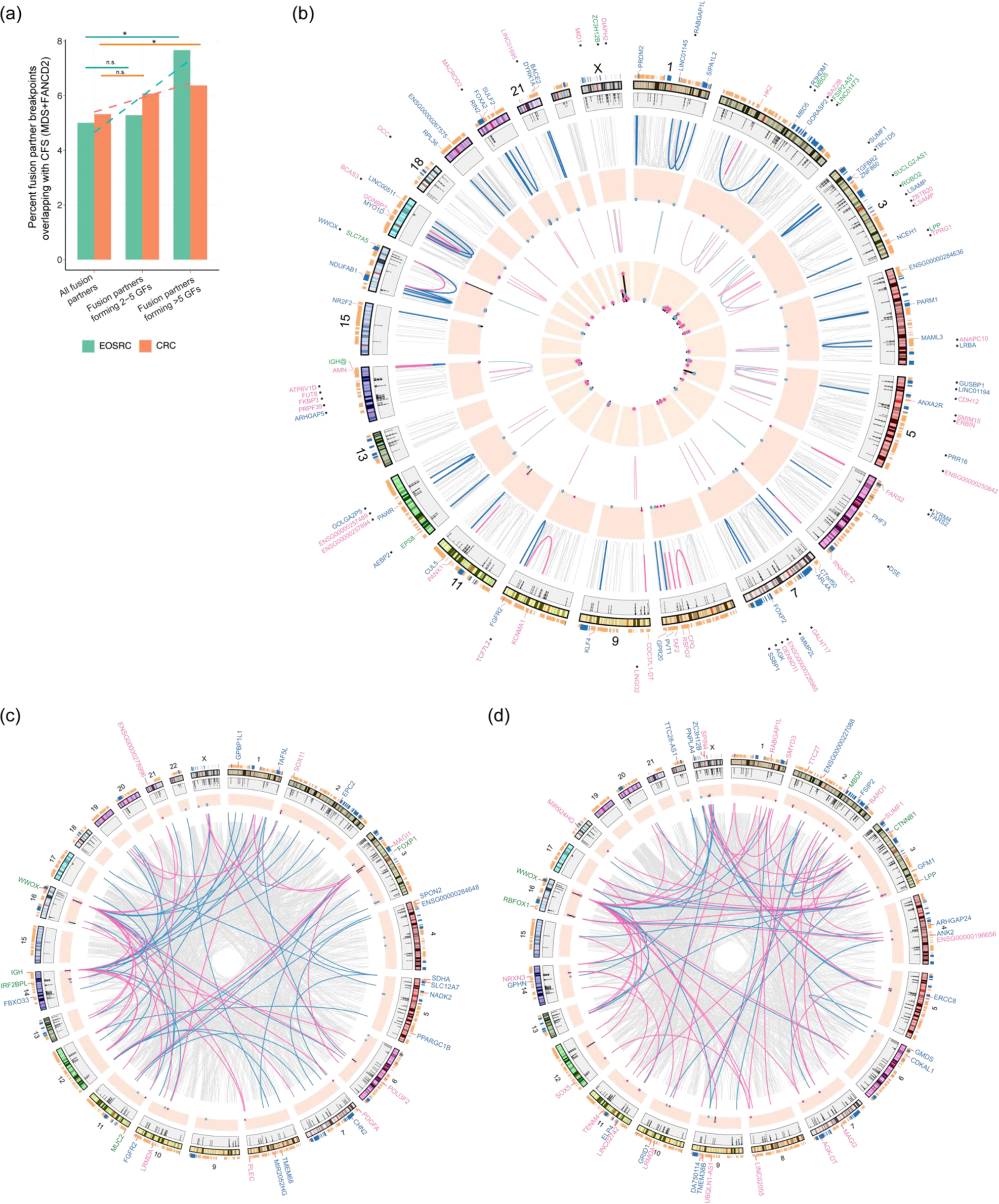
Overlap between GF breakpoints and CFS. (a) percent of fusion partner genes overlapping with CFS. Differences were considered significant at a p value < 0.05 (*P < 0.05; **P < 0.01; n.s., not significant (< 0.06)), estimated from Fisher’s exact test. (b) Circos plot showing comprehensive comparative profiling of GFs and CFSs. The track as read from the outermost to innermost is as follows – (i) average signal values from FANCD2 ChIP-Seq analysis. Blue and light orange indicate regions of high (p<0.01) and lower (p<0.05) signal intensities. (ii) Chromosomes represented as ideograms. (iii) MDS signal represented as black bars. (iv) intra-chromosomal GFs indicated with grey links. Blue and pink links indicate GFs where 5’- and 3’- gene breakpoint overlap with FANCD2 binding site, respectively. (v) Black bars indicate the sample wise recurrence of GFs involving genes whose breakpoints overlapped with FANCD2 sites. (vi) Links indicating intra-chromosomal GFs overlapping with MDS signal with same colour code as in (iv) above. (vii) Black bars indicating the sample wide recurrence of GFs involving genes with breakpoints overlapping with MDS. The outermost circle of text labels (preceded by black dots) indicate genes participating in GF formation whose breakpoints overlap with MDS sites. 5’- and 3’- partner genes are indicated in blue and pink, respectively. Genes which occur as both 5’- and 3’- partners are indicated in green. The inner circle of text labels represent the 5’ (blue) and 3’ (pink) genes whose breakpoints overlap with FANCD2 binding sites, with green indicating genes which act as both. (c, d) inter-chromosomal gene fusions with either the 5’- or 3’- gene breakpoints overlapping with FANCD2 (c) and MDS (d) sites.

Among all EOSRC fusion genes (either 5’- or 3’- fusion partners) exhibiting promiscuity whose breakpoints overlapped with CFSs (either FANCD2 or MDS), and which formed multiple intra-chromosomal GFs, the sample-wide recurrence of *SLC7A5* (26/37 samples) was the highest, followed by *ZC3H12B* (14/37), *PANX1* (6/37), *PRR16* (5/37), *FOXA2* (4/37), and *NR2F2* (3/37) (**Figure 3b, Table S4**). While chromosomes 1-2, 8, and 16-17 showed higher contribution towards intra-chromosomal GFs formed by genes overlapping with FANCD2, fusion partner genes from chromosomes 2-3, and 5 exhibited higher overlap with MDS sites. Further, from the distribution of inter-chromosomal GF breakpoints overlapping with CFS in EOSRC, GFs involving *WWOX* (17/37), *IgH* (14/37), *FOXP1* (10/37), *MAGI1* (9/37), *RBFOX1* (8/37) and *POU3F2* (4/37) were highly recurrent (**Figures 3c, d, and Table S4**). Among the GFs involving *WWOX*, the frequency of *UFM1-WWOX* was the highest (24%; 9/37) (**Figure S8c).**

Extending the analysis to TCGA-CRC revealed a similar overlap of intra-chromosomal GFs with FANCD2 and MDS sites (**Figure S9a**), but with apparent differences. Chromosomes 4-6, 8, and 15-17 contributed higher intra-chromosomal GFs formed by genes overlapping with FANCD2 sites, and several of these genes formed GFs with multiple partners across samples viz. *ZNF254* (13/157 samples), *GCC2-AS1* (13/157), *HOXB8* (8/157), *MRNIP* (7/157), *FOXA2* (6/157), *PANX1* (6/157), *PRF1* (4/157), and *DUSP6* (4/157) (**Table S4**). Similarly, chromosomes 5-8 contributed to several intra-chromosomal GFs overlapping with MDS sites of which GFs formed by *CDH12* (52/157), and *MID1* (23/157) were highly recurrent (**Figure S9a, Table S4**). Further, the inter-chromosomal GFs in TCGA-CRC overlapping with MDS sites were distributed across different chromosome pairs (**Figure S9c**). Thus, our study revealed a significant association of promiscuous GFs with CFS.

### GF formation triggers activation of transcriptionally silent genes

We next evaluated a possible correlation of higher order chromatin organization with GFs. Specific chromosomes are known to display tissue-specific spatial proximity, which could influence the frequency of inter-chromosomal translocations. We therefore analysed the relative contribution of chromosomal pairs towards GF formation. The overall contribution of intra-chromosomal events was significantly higher than inter-chromosomal events in both EOSRC and TCGA-CRC (**Figures 4a, b**). Among the intra-chromosomal events, chromosomes 1, 2, 5, 8, 7, 10, and 17 were the highest contributors towards GFs in both data sets (**Figures 4a, b**). The contribution from other chromosomes however differed between EOSRC and TCGA-CRC. We next evaluated whether chromosomes exhibiting higher tendency to form intra-chromosomal GFs also displayed higher fraction of intra-chromosomal contacts. To this end, we analysed Hi-C contact matrices generated for CRC cell lines HCT116 and SW480. A higher fraction of intra-chromosomal contacts were observed in chromosomes 1, 2, 5, 6, 7, 10, 17 (**Figure 4c**) reflecting the GF frequency trends (**Figure 4a).** To further establish the association between chromosomal contacts and GFs, we estimated chromatin contacts from Hi-C data for HCT116 (representative for EOSRC and TCGA-CRC), MCF-7 (representative for BRCA), PC3 (representative for TCGA-PRAD) and PANC-1 (representative for TCGA-PAAD). We observed a significant correlation between enriched chromosome pairs which participated in GFs in these cell lines and the corresponding chromatin contacts (**Figure 4d**), with the possible exception of PC3. Extending the analysis to GFs from tumor samples revealed a similar significant correlation between GF forming chromosome pairs and respective chromatin contacts (**Figure 4e**), suggesting that higher chromatin contacts can play a significant role in higher frequency of GFs arising from the corresponding chromosomes.

**Figure 4:**
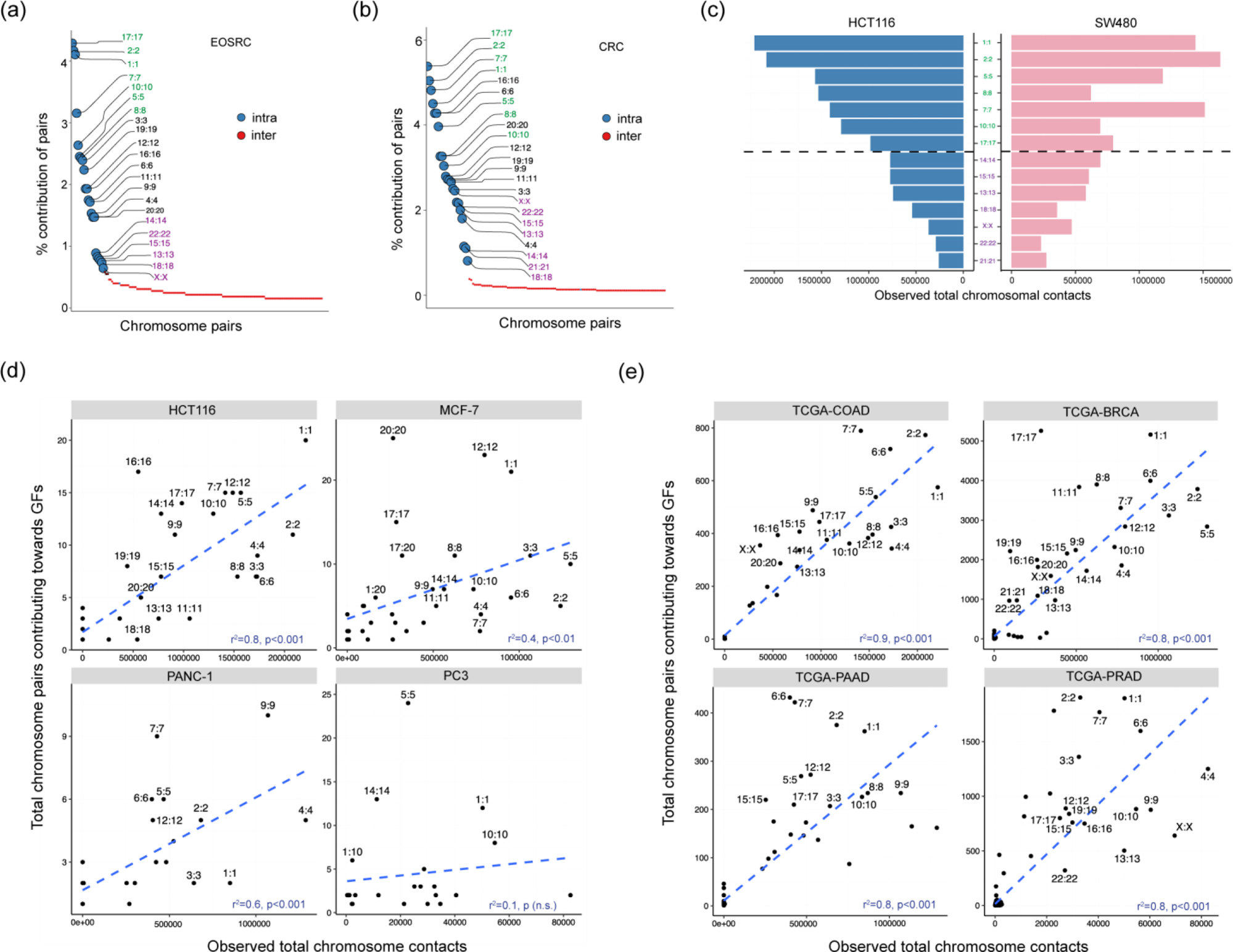
Association between chromatin architecture and GFs. (a) Contribution of chromosome pairs towards GF events in EOSRC and (b) TCGA-CRC. The chromosome pairs with highest and lowest (seven pairs from each category) contribution from intra-chromosomal events are indicated in green and purple, respectively. (c) Observed chromosome contacts from SW480 and HCT116 determined from Hi-C data. The seven highest (green) and least (purple) contributing pairs (ranked as per HCT116) are shown above and below the black horizontal broken line, respectively. (d) Association between chromosome pairs participating in GFs in HCT116, MCF-7, PANC-1, PC3 and corresponding intra and/or inter-chromosomal contacts estimated from Hi-C analysis of these cell lines. (e) Association between chromosome pairs participating in GFs in TCGA-COAD, BRCA, PAAD and PRAD and corresponding intra and/or inter-chromosomal contacts estimated from the respective cell lines. Blue dashed line indicates best fit estimated using linear regression model. r^2^ values were calculated using Pearson’s correlation estimate (P-values were estimated using Fisher’s z transformation).

Several studies have revealed chromatin to be organized into two specific compartments namely open (A) representing potential euchromatin and closed (B) representing potential heterochromatin. We estimated the compartmental organization by eigenvector decomposition of contact matrices obtained from Hi-C analysis. The estimated A compartments in HCT116, MCF-7, PC3, and PANC-1 were enriched for ATAC-Seq peaks and active histone marks, implying higher accessibility; while B compartments were enriched for repressive marks with weak or absent ATAC-Seq signals (Figure **S10a-d**). To further investigate the relation between chromatin organization and GFs, we analysed the concordance between chromatin compartment assignments and corresponding expression levels of fusion partner genes for all GFs detected in cell lines. The assigned chromosome compartments showed concordance with the expression levels of all fusion partner genes in the four cell lines i.e. the fusion partner genes which were assigned to A compartment showed higher expression compared to those assigned to the B compartment (**Figure S11**). These observations validated the accuracy of chromatin compartment assignment to the GF partner genes.

We then proceeded to determine the association of GFs with chromatin organization. It was observed that a higher fraction of A (compared to B) compartment genes participated in fusion formation in the cell lines (with the exception of PC3) (**Figure S12a**). This was expected, given that open chromatin is gene rich and exhibits active transcription and is therefore vulnerable to replication and transcription related stresses as well as exposure to external insults. In contrast, close chromatin is gene poor and transcriptionally silent. Further, in a majority of GF pairs, both the partner genes were associated with A chromatin compartment (A-A GFs) in all four cell lines (**Figure S12b**). We next hypothesized that fusion of a 5’- gene located in the transcriptionally active A compartment with a 3’- gene located in the transcriptionally silent B compartment, could potentially activate the expression of the otherwise silent 3’- partner. Indeed, the expression levels of B compartment genes participating in A-B GFs were consistently higher than those not involved in GF formation in all four cell lines (**Figure S12c**). In contrast, the expression levels of 3’ partners in A-A GFs were elevated to a lesser extent **(Figure S12d)**.

We next attempted to validate these novel observations in tumor samples. We first verified that the assigned chromosome compartments of all fusion partner genes for GFs detected in tumor samples showed concordance with the expected expression levels in the corresponding cell lines (**Figure S13a-d**) as well as in the tumor samples themselves (**Figure S13e-f).** Similar to the results obtained in cell lines, a higher fraction of A compartment genes participated in fusion formation compared to fraction of B compartment genes (**Figure S13g).** Further, A-A GFs were observed at a higher frequency than other combinations across all tumor datasets **(Figure S13h**).

In order to confirm modulation of expression levels of 3’- fusion partners in A-B GFs by the 5’-partners, we used the EOSRC, TCGA-CRC, BRCA, PAAD and PRAD RNA-Seq data sets to estimate the transcript abundances of all genes involved in GF formation. The median expression levels of 3’-partner genes in A-A GFs were elevated in samples with GF compared to samples without them in EOSRC as well as in all TCGA cancer datasets (**Figures 5a, S14)**. More importantly, the median expression levels of the 3’-partner genes were significantly elevated in samples with A-B GFs compared to samples without GF in all tumor datasets (**Figures 5b, c**). A quantification of the fraction of 3’-partner genes which exhibited a significant increase in their transcript levels revealed that in EOSRC, of all the 3’- partner genes from A-B GFs (total – 289), 21.4% showed a significant increase (fold change (fc) >= 1.5) in their median expression levels in samples with GFs compared to samples without (p-value<0.01, Wilcox test). This fraction was significantly higher than the 3’- partners from A-A GFs, where it was just 10% (p < 0.05, Wilcox test) (**Figures 5d, e**). In TCGA-CRC, 19% 3’-partners from A-B GFs had an fc >= 1.5 (p < 0.01, Wilcox test), which was again significantly higher than 3’-partners from A-A GFs (7%, p < 0.01, Wilcox test) (**Figures 5d, e**). While in TCGA-BRCA, 20% A-B GF 3’-partners exhibited fc>=1.5, as against 11% 3’-partners from A-A (p<0.01, Wilcox test). This fraction was 14% and 7% (A-B vs A-A 3’-partners, fc>=1.5, p<0.01, Wilcox test) for TCGA-PAAD. The difference in the fraction of 3’-partners was least for TCGA-PRAD where it was 8% (A-B GFs) and 6% (A-A GFs) (p-value < 0.05, Wilcox test) (**Figures 5d, e**). These observations validated our initial hypothesis that a significant fraction of 3’- partner gene in A-B GFs could display enhanced expression due to GF formation. Interestingly, the 21.4% 3’- partners participating in A-B GFs in EOSRC with fc > 1.5 exhibited an enrichment of pathways including coagulation, hypoxia, and complement system while similar 3’ partner genes in TCGA-CRC exhibited moderate enrichment in pathways such as G2-M checkpoint and E2F (**Figure 5f**).

**Figure 5:**
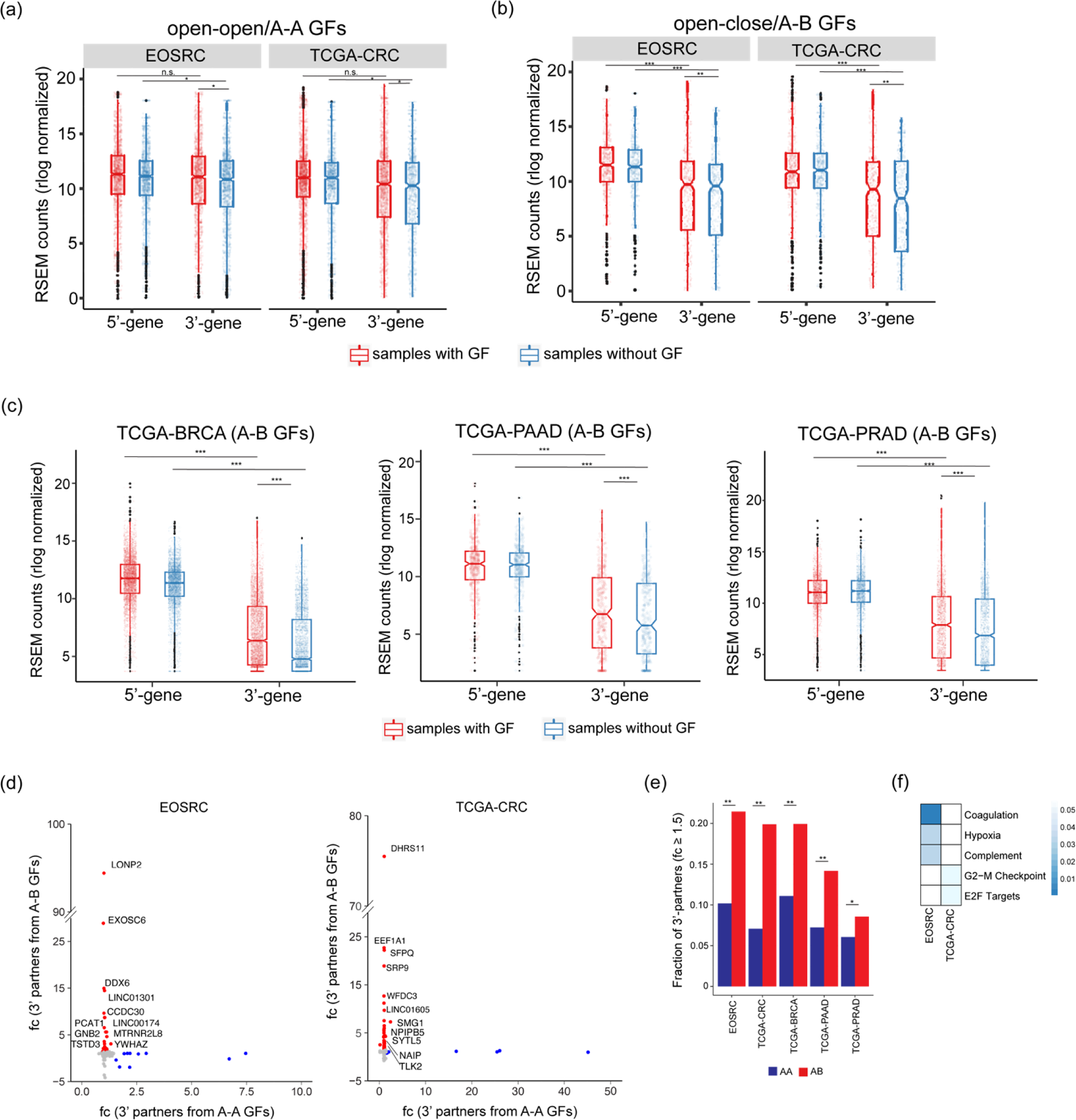
(a-c) Transcript levels of all 5’- and 3’- partner genes in samples with and without GFs in (a) A-A (open-open) category and (b) A-B (open-close) category in EOSRC and TCGA-CRC, (c) in A-B category in TCGA-BRCA, PAAD and PRAD. P-values (Wilcoxon rank test): n.s., not significant; *, < 0.05; **, < 0.01; ***, < 0.001. (d) Fold change (fc) values for 3’-partner genes in A-A vs A-B GFs in EOSRC and TCGA-CRC. 3’-partners with fc ≥ 1.5 are highlighted in red (A-B GFs) and blue (A-A GFs). (e) Fraction of 3’-partners in A-A and A-B GFs with fc ≥ 1.5. P-values: *<0.05, ** <0.01. (f) MSigDB hallmark pathways associated with 3’- partner genes exhibiting increased expression (fc≥1.5) in A-B GFs.

The elevation in the transcript levels of 3’ partner genes participating in A-B GFs in EOSRC tumors (vs tumors that did not exhibit the GFs) was validated from the analysis of RNA-seq data (**Figures 6, S15**). Interestingly, the 3’ partner genes located in B compartment exhibited insignificant transcript levels in all samples except in the sample where it formed fusion with 5’ partner from A compartment (**Figure 6**); this also validated the accuracy of chromatin compartment assignment based on Hi-C data. The only exception was *RSPO3* which exhibited elevated expression even in samples where it was not participating in A-B GF (**Figure 6**). This could point towards other modes of *RSPO3* activation in the absence of GFs^63^. In contrast, the transcript levels of the 5’ partners in the same GFs did not exhibit any such trend (**Figure 6**). We further confirmed the elevated expression of 3’ partner genes from select A-B GFs in EOSRC samples by RT-qPCR, while the levels of 5’- partners varied between the samples (**Figure 6**). The elevated expression of 3’ partner gene in A-B GFs was also validated from the transcript levels estimated from RNA-Seq analysis of TCGA-CRC (**Figure S16**), BRCA, PRAD and PAAD. (**Figure S17**). These results conclusively support our inference that in GFs formed between genes from orthogonal compartments (A-B GFs), the expression of the 3’- partner genes are elevated significantly compared to when the same gene is not participating in A-B GFs.

**Figure 6:**
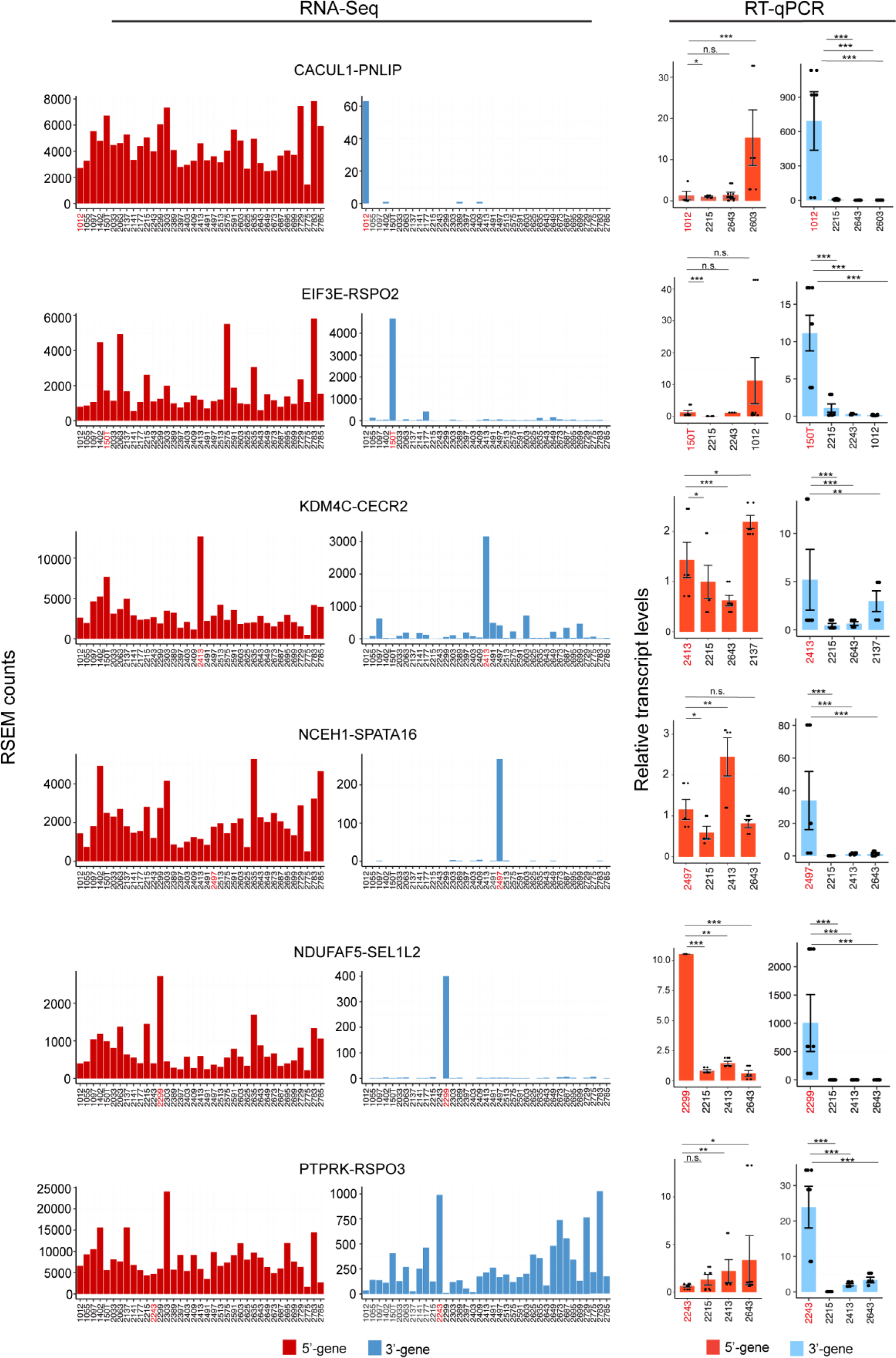
Sample-wise transcript levels of 5’- and 3’- partner genes for select A-B GFs estimated from RNA-Seq data generated for EOSRC and RT-qPCR based validation of transcript levels for respective 3’-partner genes. Sample with the GF is indicated in red colour on the x-axis. P-values were calculated using Wilcoxon rank test. n.s., not significant; *, < 0.05; **, < 0.01; ***, < 0.001.

Finally, we determined the fraction of A or B compartment genes forming GFs that exhibited association with CFS. Interestingly, a higher fraction of GF partners (whether 5’ or 3’) localizing to the B compartment exhibited overlap with CFSs, compared to fusion partners from the A compartment, in both EOSRC and TCGA-CRC (**Figure S18a**). Further, a similar analysis of A-A, A-B, B-A, and B-B GFs revealed that a higher fraction of B (compared to A) compartment genes participating in GFs overlap with CFS loci (**Figure S18b**) in both EOSRC, and TCGA-CRC. Association of a larger fraction of fusion forming B compartment genes with CFS points to an important role played by CFS in driving GF events with relevance to tumorigenesis, given the ‘inert’ nature of B compartment genes. Overall, these results indicate that chromatin organization and genome stability show an intricate and complex interplay leading to promiscuous and recurrent GFs, while playing a significant role in activating expression of B compartment genes.

## Discussion

With rapid advances in deep sequencing technologies and computational approaches for data analysis, exploratory studies on genome wide *de novo* GF identification has become increasingly feasible. While, CRC tumorigenesis has been well studied in terms of recurrent somatic mutations, altered DNA copy number and deregulated gene expression, somatic genome rearrangements have been less well examined. Furthermore, knowledge of GFs in EOSRC, a poorly understood albeit clinically important CRC subtype, is absent. Here, we proceeded to comprehensively characterize GFs in EOSRC and perform a comparative analysis of GFs identified in conventional CRC and other non-CRC cancer types. We identified a higher frequency of per sample GFs in EOSRC compared to CRC data from TCGA, owing probably to the greater sequencing depth of the data generated for the former. We also retained GFs which involved (non-coding) ncRNAs (including long ncRNAs, putative circular RNAs and miRNAs) as fusion partners in our study, which could be another reason for higher frequency of GF detection in EOSRC. Several previously well documented CRC GFs were recapitulated in this study. More importantly, we discovered recurrent, albeit previously unidentified GFs in both EOSRC and TCGA-CRC. A notable example is the *CNTRL-PCID2* fusion identified in EOSRC. *PCID2* (encoding PCI domain containing 2) amplification caused *PML* (promyelocytic leukemia) degradation, which resulted in increased cell growth and migration due to enhanced canonical Wnt/β-Catenin signalling in CRC^64^. Additionally in EOSRC, several novel GFs such as *KDM4C-CECR2*, *LGALS9-KIAA0100*, and *NDUFAF5-SEL1L2* involving functionally important and potentially actionable genes were identified and validated using RT-PCR and Sanger sequencing.

Despite the steady progress in the development of whole genome and transcriptome profiling methods, bioinformatics pipelines for GF detection are still under active development. Due to these limitations, our study is expected to have missed potential complex or bridged fusion events involving more than two fusion partners. Additionally, large scale validation of GFs remains challenging, with overall validation rates by RT-PCR varying between different tumor types. Of note, the EOSRC GFs confirmed through RT-PCR and Sanger sequencing in this study represent the largest number validated in a single study, to the best of our knowledge.

We evaluated a GF network arising from the promiscuity in genes that participated in multiple GFs. Earlier reports have suggested this promiscuity leads to a highly divergent and interconnected network, with the potential to link and rewire complementary and/or diverse biological pathways^59^, similar to our findings. It may be pertinent to explore this aspect in greater detail in CRC and additional cancer types. The observation of higher degrees exhibited by Immunoglobulin (Ig) loci (*IgH*, and *IgK*) was particularly interesting as the significance of these loci in GFs have been well studied in liquid cancers including multiple myeloma^65^, where the frequency of Ig fusions is found to be highest. Since Ig loci can contribute multiple regulatory elements^66^, it can be speculated that they can potentially drive expression of their fusion partners. In this study, multiple oncogenes e.g. *CDK6, CALR, TAF15*, and *CRLF2* were observed to form GFs with *IgH* in EOSRC. Similarly, *EEF1A1* shows high transcript levels in a range of cell types^67^; formation of multiple fusions involving this gene as a 5’ partner provides additional mechanism of altering expression levels of the 3’ partner genes. This becomes relevant for tumorigenesis especially if the 3’ partner is oncogenic; we identified GFs involving *EEF1A1* and 3’ oncogenic partners such as *CDH17* and *TMSB4X*.

CFSs are more likely to ‘break’ during tumor formation due to replication stress and breakpoints involved in oncogenic amplification have been previously linked to CFS loci^68^. Moreover, genes lying within these loci are more likely to form fusions with multiple partners. However, a direct association between CFS and GF breakpoints was not comprehensively explored earlier. We have demonstrated this association in EOSRC and TCGA-CRC by correlating promiscuous GF breakpoints with CFS loci. It will be interesting to extend observations to other tumor types, especially those known to exhibit chromosomal instability like lung (squamous and adenocarcinoma) and breast adenocarcinoma.

Studies in cell lines derived from liquid cancers have shed light on the spatial proximity of GF partner genes^28^, influenced partly by co-localization of specific chromosome pairs, which may differ between tissue types^69^. Our data on chromosomal contacts indicate that GFs from solid tumors may also exhibit a similar association. We are currently analysing high resolution Hi-C data to assess whether close proximity of chromatin segments facilitates GF formation by correlating anchor points of chromatin loops with GF breakpoints. We further performed a parallel analysis of gene fusions (from transcriptome data) and chromatin compartments (from Hi-C data), which enabled us to tease out relative contributions of A and B compartments towards GF formation. Though a higher proportion of A (compared to B) compartment genes contributing to GF formation was expected, ours is the first documentation of such an association. Analysis of chromatin compartment associated with GF partners led to the discovery of a novel mode of 3’- gene activation in cancer resulting from the fusion between orthogonally located 5’ (A compartment) and 3’ (B compartment) partner genes. Fusion formation appears to drive the expression of the 3’ partner gene in A-A as well as A-B GFs but the extent of activation is higher in the latter, perhaps given the inherent silencing of the genes located in the B chromatin compartment. Although, the activation of 3’-partners due to GF formation has been demonstrated earlier^70^, this is the first report of alteration of expression levels of 3’ partner genes based on their chromatin compartment localization. It will be interesting to further study the possible mechanism of activation of the 3’ partner genes in A-B fusions. Do the genes remain associated with their respective compartments even after the fusion? Or does the 3’ partner originally associated with the B compartment translocate to A compartment following the fusion, thus facilitating its activation? Similarly, do the A-B or B-A fusions occur between genes that are located at the transition between the two compartments?

The significantly higher expression of the otherwise silent 3’-partner genes in A-B GFs in tumors (**Figure 7**) gain significance especially if the 3’- partner gene has oncogenic functionality. We identified A-B GFs where expression levels of B compartment genes such as *RSPO3* (*PTPRK-RSPO3*), *RSPO2* (*EIF3E-RSPO2*), and *CECR2* (*KDM4C-CECR2*) were significantly elevated. Both *RSPO2* and *RSPO3* have been shown to be potential oncogenic therapeutic candidates in CRC harboring Wnt pathway mutations^63^. RSPO2 has also been proposed as oncogenic driver in invasive breast cancer^71^ as well as prognostic markers in prostate^72^ and ovarian^73^ cancers. Another interesting example of a B compartment gene exhibiting elevated expression in an A-B GF (in TCGA-CRC) is *PDIA3 (ACTB-PDIA3)*. The *PDIA3* (protein disulphide isomerase A3) gene has been described as a potential proto-oncogene^74^ and its upregulation was linked to higher CRC cell survival and resistance to radio/chemotherapy in CRC tissues^75^. Of note, most other 3’ partner genes exhibiting this novel mode of activation identified in EOSRC and TCGA-CRC are not studied in the CRC context. For example, *CECR2 (KDM4C-CECR2)* is a bromodomain containing protein involved in chromatin remodelling^76^, and has been shown to be recruited by *RELA* (v-rel avian reticuloendotheliosis viral oncogene homolog A), to increase chromatin accessibility and activation of metastasis promoting target genes such as *MMP2* and *VEGFA*^77^ in metastatic breast cancer. It will be interesting to evaluate role of these potential novel CRC genes.

**Figure 7:**
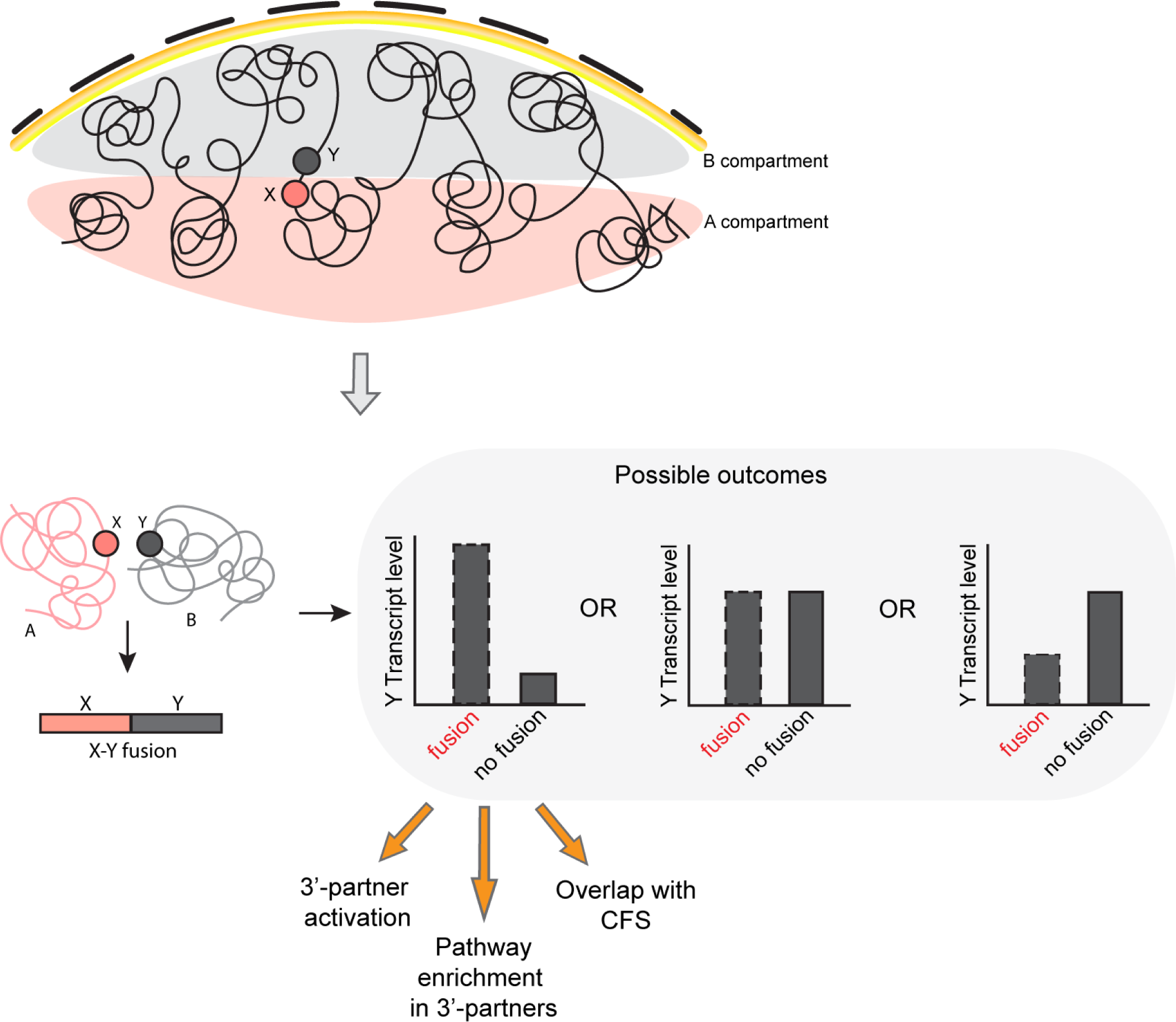
Schematic representing possible mode of activation of 3’ partner gene in A-B GFs.

We are currently evaluating the possibility of potential trans-splicing (as against DNA level) events leading to GF formation. Given the widely accepted concept of ‘transcription factories’ where different chromosomes converge at a highly localized area within the nucleus to facilitate enhancer mediated activation of multiple genes^78^, trans-splicing events leading to GF formation could be envisaged.

EOSRC remains a poorly understood albeit aggressive CRC subtype despite the recent significant increase in worldwide incidence^79^, underscoring the importance of defining novel therapeutic options. It would be worthwhile to evaluate potential therapeutic utility of the novel and/or recurrent and potentially actionable GFs reported in this study, especially involving kinases (*AAK1-GFPT1, MUC2-PLK1*). Further, translation of GF transcripts (for in-frame fusions) is expected to give rise to chimeric proteins and such neo-antigens can be potentially utilized for diagnostic and prognostic utility (given their relative absence in normal tissue) besides being potentially actionable^80^. In addition, such chimeric proteins can be expected to be processed and presented with MHC molecules; a facet that can be potentially harnessed for therapeutic options involving ‘killer’ immune cells.

In conclusion, the current study presents a comprehensive landscape of GFs in EOSRC (and TCGA-CRC), highlighting a critical set of novel and recurrent fusions of potential clinically utility. The analyses also revealed a significant overlap between promiscuous fusion partners and well-defined CFS loci. More importantly, we provide the first evidence of activation of transcriptionally silent B compartment genes owing to their fusion with genes from orthogonal A chromatin compartment, a novel discovery which was validated in several cancer types. Our study has therefore opened up several new aspects for future investigations in CRC (and potentially in other cancer types).

## Acknowledgements

The authors acknowledge Mandla Vasanth Kumar for his assistance in carrying out RT-PCR. A.G. received funding from the Women Scientist Scheme – A, Department of Science and Technology (DST-WOSA), Government of India and the National Postdoctoral Fellowship (NPDF) scheme, Science and Engineering Research Board (SERB), Department of Science and Technology, Government of India. The authors thank Dr Pratyusha Bala for designing the RNA-Seq experiments and the Sophisticated Equipment Facility (SEF), CDFD, for Sanger Sequencing. The authors thank Dr Sara Anisa George, Laboratory of Molecular Oncology, Centre for DNA Fingerprinting and Diagnostics, Hyderabad, India, Dr Jaya S Tyagi, Department of Biotechnology, All India Institute of Medical Sciences, N. Delhi, India and Dr J Gowrishankar, Indian Institute of Science, Education and Research, Mohali, India, for critical reading of the manuscript.

## Author Contributions

M.D.B. and A.G. conceived the study; A.G. performed the computational analyses; A.G. and S.A. performed RT-PCR and RT-qPCR assays. A.G. and M.D.B. compiled and analysed the data. A.G. and M.D.B. wrote and revised the manuscript.

## Data availability

All the raw sequencing data generated in this study has been submitted to the GEO database with accession ID GSE253106. The list of all primers used has been provided in table S5. Custom R codes used for generating figures can be accessed from the Github repository (https://github.com/asmitagpta/crc_genomics).

